# A minimal robophysical model of quadriflagellate self-propulsion

**DOI:** 10.1101/2021.03.14.434582

**Authors:** Kelimar Diaz, Tommie L. Robinson, Yasemin Ozkan Aydin, Enes Aydin, Daniel I. Goldman, Kirsty Y. Wan

**Affiliations:** School of Physics, Georgia Institute of Technology, Atlanta, GA 30332, United States of America; Living Systems Institute & College of Engineering, Mathematics, and Physical Sciences, University of Exeter, EX4 4QD, United Kingdom

**Keywords:** swimming, cell motility, robot, bio-inspired, gait, quadrupedal locomotion, algae

## Abstract

Locomotion at the microscale is remarkably sophisticated. Microorganisms have evolved diverse strategies to move within highly viscous environments, using deformable, propulsion-generating appendages such as cilia and flagella to drive helical or undulatory motion. In single-celled algae, these appendages can be arranged in different ways around an approximately 10 *µ*m cell body, and coordinated in distinct temporal patterns. Inspired by the observation that some quadriflagellates (bearing four flagella) have an outwardly similar morphology and flagellar beat pattern, yet swim at different speeds, this study seeks to determine whether variations in swimming performance could arise solely from differences in swimming gait. Robotics approaches are particularly suited to such investigations, where the phase relationships between appendages can be readily manipulated. Here, we developed autonomous, algae-inspired robophysical models that can self-propel in a viscous fluid. These macroscopic robots (length and width = 8.5 cm, height = 2 cm) have four independently actuated ‘flagella’ that oscillate back and forth under low-Reynolds number conditions (Re *∼ 𝒪*(10^*−*1^)). We tested the swimming performance of these robot models with appendages arranged in one of two distinct configurations, and coordinated in one of three distinct gaits. The gaits, namely the pronk, the trot, and the gallop, correspond to gaits adopted by distinct microalgal species. When the appendages are inserted perpendicularly around a central ‘body’, the robot achieved a net performance of 0.15 *−* 0.63 body lengths per cycle, with the trot gait being the fastest. Robotic swimming performance was found to be comparable to that of the algal microswimmers across all gaits. By creating a minimal robot that can successfully reproduce cilia-inspired drag-based swimming, our work paves the way for the design of next-generation devices that have the capacity to autonomously navigate aqueous environments.

## 1. Introduction

The capacity for self-generated movement is a distinguishing feature of most living organisms. In the macroscopic world, locomotion is typically associated with inertia [1]. On the other hand, movement at the microscopic scale is subject to low Reynolds number physics, and cannot take advantage of inertial coasting. Without motility, a bacterium can only coast a minuscule distance an order of magnitude below the Ångström scale [2]. Over billions of years of evolution, microorganisms have become adept at swimming, evolving distinct mechanisms for powering and maintaining self-movement through a fluid, often achieving speeds of several tens of body lengths per second. This active motility confers a significant survival advantage, allowing microbes to navigate freely towards regions or locations where nutrients or resources are more plentiful [3]. Depending on the arrangement and number of locomotor appendages, single cells can execute swimming gaits that are surprisingly reminiscent of animals. For example, the model biflagellate alga *Chlamydomonas* actuates two equal-length flagella in a breaststroke [4], while quadriflagellate algae (single cells with four-flagella) exhibit distinctive quadrupedal gaits such as the trot or the gallop [5] (Fig. 1A,B).

**Figure 1.**
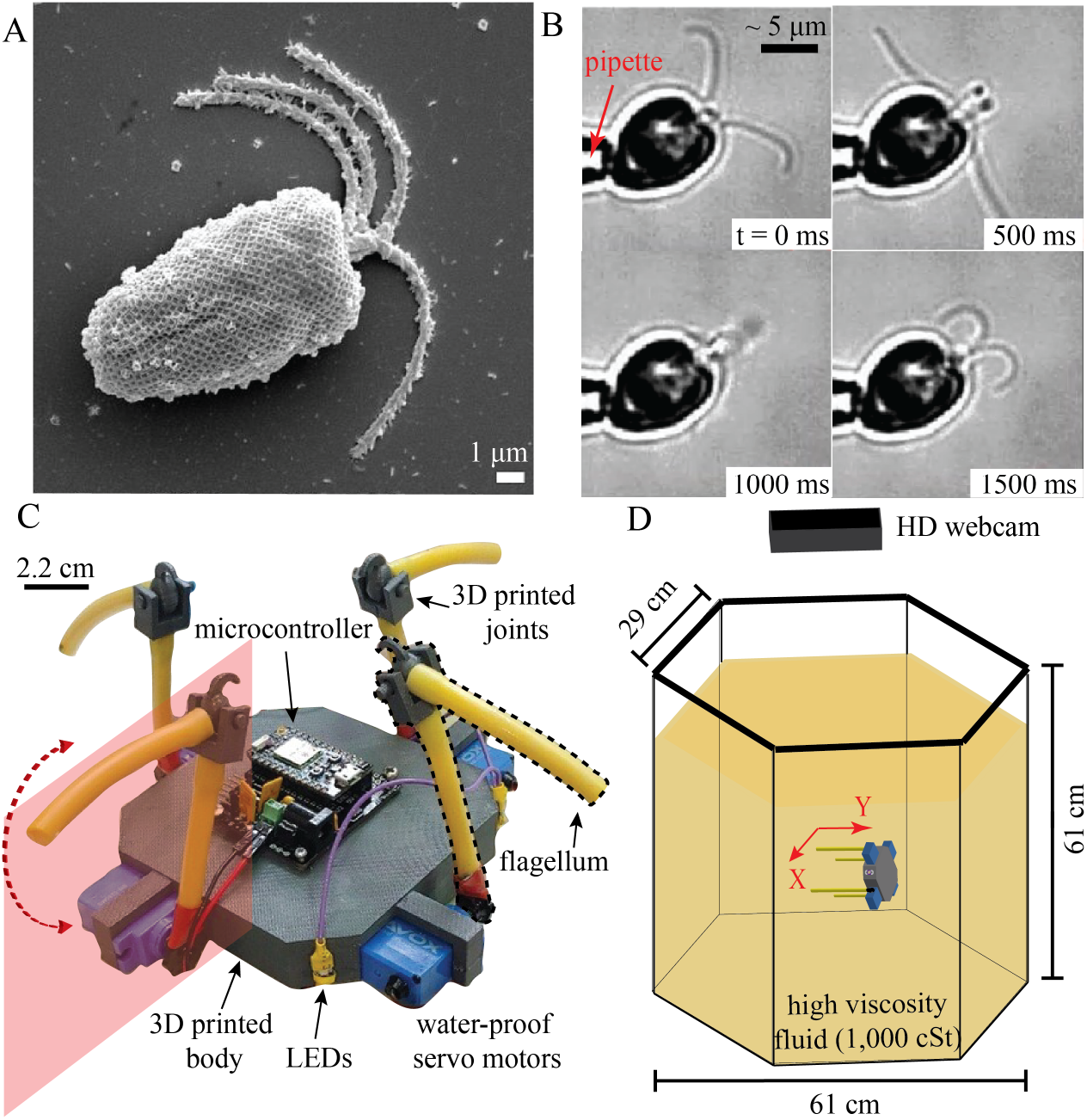
Design and fabrication of a dynamically-scaled robophysical model of a microswimmer with four flagella. (A) Scanning electron microscope (SEM) image of *Pyramimonas gelidicola* [36]. (B) Snapshots of *Pyramimonas parkeae* held by a pipette. (C) Robophysical model of quadriflagellate algae. (D) Experimental set-up. The arena is a hexagonal tank filled with mineral oil of high viscosity, to match the low Reynolds number regime experienced by the algae.

In recent years, advances have been made in understanding the biomechanics of microswimming. Here, the Reynolds number is small, *Re* = *UL/ν*, where *L* is a typical lengthscale of the swimmer, *U* a typical velocity scale, and *ν* is the kinematic viscosity of the fluid. Equally important is the oscillatory Reynolds number *Re*^osc^ = *L*^2^*ω/ν* [6], where *ω* the typical stroke frequency (which sets a tip velocity of *ωL*). When both are small, flows are then governed by the Stokes equations: 0 = ∇*p* − *µ* ∇^2^**v** and ∇. **v** = 0 (where **v** and *p* are the flow and pressure fields), and have no explicit time-dependence. Microorganisms are able to break time-reversal symmetry using non-reciprocal strokes or body deformations, often involving whip-like appendages called cilia and flagella [2, 7]. While bacteria make use of rigid helical flagella [8], eukaryotes actuate motile cilia which produce asymmetric waves of propulsion [9, 10]. For a microorganism oscillating a 10 *µm* flagellum at 50 Hz, *Re ∼*10^*−*3^, and *Re*^osc^ *∼*10^*−*2^. One further asymmetry is required for forward propulsion [11] - this is shape asymmetry, which is ensured by the slender aspect ratio of all cilia and flagella (about 100). A rod sweeping through a fluid in the direction perpendicular to the axis of the rod experiences approximately twice the drag compared to when it is moved in the parallel direction [12]. Organisms across all scales have been found to exploit this basic anisotropy in frictional forces for locomotion [13, 14, 15].

Despite the adoption of cilia and flagella as a common propulsion mechanism, the microscale locomotion strategies of microorganisms have diversified significantly across different phyla [16]. It is not well-understood why different gaits exist nor how they are coordinated. For centuries, locomotor gaits have been studied in the context of terrestrial animals, where the sequences of relative movement sustained by subsets of limbs or legs have fascinated researchers. In vertebrates, gaits are thought to be generated by central pattern generators (CPGs) [17]. But how can orderly, deterministic appendage coordination occur in single cells in the absence of nervous control [16, 18]? Recent theoretical and experimental work have show that dynamic gait selection, at least in flagellates, appears to be an active and species-dependent process driven by intracellular and mechanical coupling [18, 19]. Notably, distinct quadriflagellates can self-propel at different speeds despite an apparently identical arrangement of flagella around the cell body [5, 20]. Since the ancestral form of the green algal lineage may have have been a unicell with four flagella [21], there is much incentive to understand the precise mechanisms of appendage coordination in such systems.

In the quest to address these open questions of movement control, extant organisms can provide only a limited parameter space of possibilities in terms of size, shape, beat frequency etc, often making it challenging to investigate certain configurations or physical regimes. Theoretical and computational approaches have been instrumental in shaping our understanding of active propulsion [12, 22], but these can be computationally expensive or reliant on simplifying assumptions. Meanwhile robophysical modelling has emerged as a powerful and versatile technique for elucidating organismal behaviour by engineering customised configurations that can be easily tested in controlled laboratory settings [23, 24, 25]. The revolution in robophysical modelling has been driven in part by cheap electronics (motors, microcontrollers), and increasingly accessible control technologies that can complement theoretical modelling to provide real biological insights [24, 26]. However, trying to model cell movement is a significant conceptual challenge when working *at the microscale*. Even though increasingly controllable micro- and nano-devices have been fabricated to mimic the locomotive behaviours of biological swimmers [27, 28], these are overwhelmingly driven by external magnetic, electric or chemical fields. For instance magnetic fields are often unable to deliver the fine spatial control, required to independently actuate individual artificial cilia in a given array or network. Thus detailed investigations of the effect of gait on microswimming has mostly been restricted to theoretical microswimmers [29, 30], occasionally in artificial or colloidal swimmers [31], but seldom in microrobots [32].

The intrinsic limits of device manufacture at small scales severely undermines the suitability of microbots as realistic models of cell motility. To understand the influence of gait on self-propulsion at low-Reynolds number, our goal is to build a dynamically-scaled robophysical model which is truly *self-powered*, where the movement of individual locomotor appendages can be prescribed and controlled independently. In contrast to traditional ‘microrobots’, the large size allows us to explore and take advantage of increasingly sophisticated electronics and control architectures [33, 34]. We can readily reprogram these “roboflagellates” to execute specific swimming gaits, making them uniquely suited to testing theories of bio-inspired and autonomous locomotion at low-Reynolds number. This paper is organised as follows. We first identified and measured the relative swimming performance of three species of quadriflagellate algae that exhibit near-identical morphology but distinct swimming speeds. Next we built a centimetre-sized robot which can self-propel in high-viscosity fluid when mimicking the asymmetric beat pattern of the algal flagella, verifying that low-Reynolds number kinematics are recapitulated. By arranging the robotic flagella in one of two possible configurations (parallel or perpendicular) relative to a central “cell body”, we imposed and tested three distinct flagellar actuation patterns (gaits) that occur naturally in the algal flagellates, namely the pronk, the trot, and the gallop. In each case, we compared the hydrodynamic swimming performance of the robot to that of the corresponding algal species. Finally, we discuss the relevance of these results for understanding how functional differences in swimming performance may arise from morphologically similar structures, and highlight the implications of this from an eco-evolutionary perspective.

## 2. Methods

### 2.1. Microalgal culturing and imaging

Three species of algae (*P. parkeae, P. tetrarhynchus* and *C. carteria*) were cultured axenically according to previously published protocols [5, 18]. Free-swimming individuals were tracked in open microfluidic chambers using a high-speed camera (Phantom Vision Research). Brightfield imaging was conducted with 40x or 60x objectives using standard inverted microscopes (Leica DMi8 and Nikon T2000-U) under white light illumination. Free-swimming trajectories were obtained from high-speed videos in which single cells crossed the focal plane, with the use of the open source software TrackMate (Fiji) [35]. Ten cells per species were used to determine the performance of each swimming gait (Supplementary Video 1). Tracks in which cells performed transient gaits, tumbles, or changed directions were not used in this analysis. The body length of each cell was measured along the long axis (anterior-posterior) of the organism. An average body length of 13.95 *±* 2.05 *µ*m, 12.54 *±* 0.65 *µ*m, and 12.82 *±* 0.72 *µ*m was found for *P. parkeae, P. tetrarhynchus* and *C. carteria* respectively.

### 2.2. A self-powered roboflagellate

We designed a dynamically-scaled robot to ensure that the robotic model can be self-powered and does not require external fields - all controllers and servos are fully self-contained (Fig.1C). We performed robotic experiments in a highly viscous fluid (mineral oil, McMaster, 1000 cSt, product no. 1401K75) to match the low Reynolds number regime experienced by the algae (Figure 1C). A subset of trials were conducted in glycerin (vegetable glycerin, Blue Water Chem Group, product no. B07FQWDTH7) of comparable viscosity to the mineral oil, to enable better visualisation and tracking of appendage movement. Each robot consisted of a 3D printed body (length and width = 8.5 cm, height = 2 cm) attached to four flagella that are independently actuated by waterproof servo motors (Savox, product no. SW0250MG). Each appendage was oriented such that the stroke lies in the plane perpendicular to the body (Figure 1D). Foam (FOAMULAR Insulating Sheathing (IS) XPS Insulation) was attached on the robot body to achieve neutral buoyancy and allow it to swim untethered. Commanded appendage positions were achieved using a micro controller (Photon, Particle, part ID: PHOTONH) that allowed us to actuate our robot with the use of Wi-Fi. Our micro-controller and each motor were connected via a IOT Servo Shield (Actuonix, part ID: IOT-SHIELD-PHOTON), a circuit board specific to our micro controller. Four LEDs were placed on the 3D printed body to facilitate tracking. The robot was powered with three lithium ion polymer batteries (3.7 V, 2500 mAh), each powering directly the micro controller, the motors, and any attached LEDs. With this micro controller, the robots were able to achieve self-propulsion.

### 2.3. Actuation of robotic flagella

Each robotic appendage comprised a two-link flagellum (length = 6.5 cm, diameter = 3.1 mm, polypropylene-based thermoplastic elastomer (TPE)) connected by a 3D printed joint allowing each flagellum to passively bend and break drag symmetry (Figure 1C). Inspired by the flagellar beating waveform of the organisms, we implemented a simple two-link flagellum in the robot that was able to break time-reversal symmetry (Figure 2). An irreversible stroke pattern was achieved with the use of 3D printed hinges between the two flagella segments. Instead of actively prescribing the shape of the flagella over a beat cycle, symmetry breaking was achieved passively. No external control such as magnetic fields were used, our robot was completely open loop. Each gait maintained a constant phase difference between adjacent flagella set by prescribed joint angles of the proximal segment (Figure 3, Supplementary Video 2). Each gait was uploaded to the microcontroller via Wi-Fi, allowing the controllers to actuate the motors offline. Unless otherwise specified, all gaits were prescribed with a flagellar beat frequency of 0.14 Hz. The applied torque was constant throughout each beat cycle (3.5 kg/0.34 Nm operating at 4.8 V).

**Figure 2.**
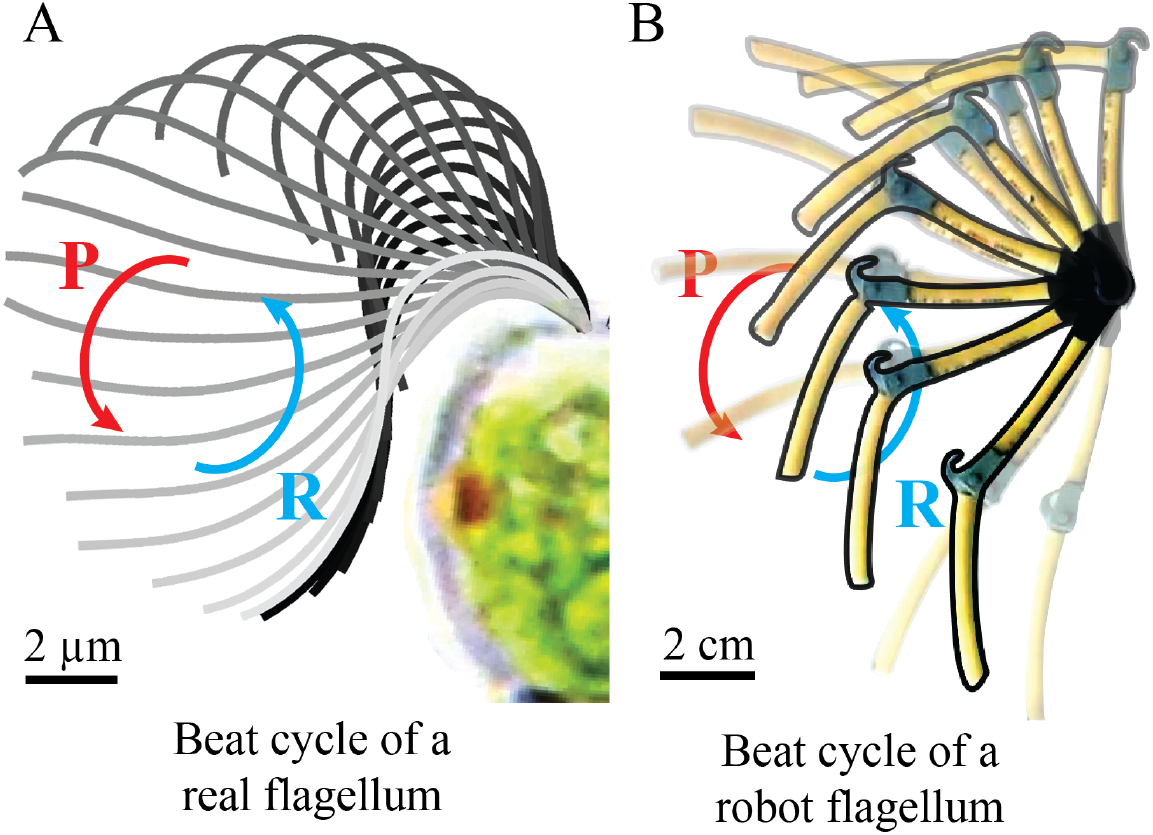
Breaking time-reversal symmetry with a hinged two-link bio-inspired flagellum. One beat cycle of an (A) algal flagellum compared to a (B) robot flagellum. P: power stroke, R: recovery stroke. Each robot flagellum segment has a length of 6.5 cm and diameter of 3.1 mm. Asymmetric beat patterns are achieved via a 3D printed joint. The movement patterns of the algal flagellum was measured in water, for the robot this was visualized in a high-viscosity fluid (glycerin).

**Figure 3.**
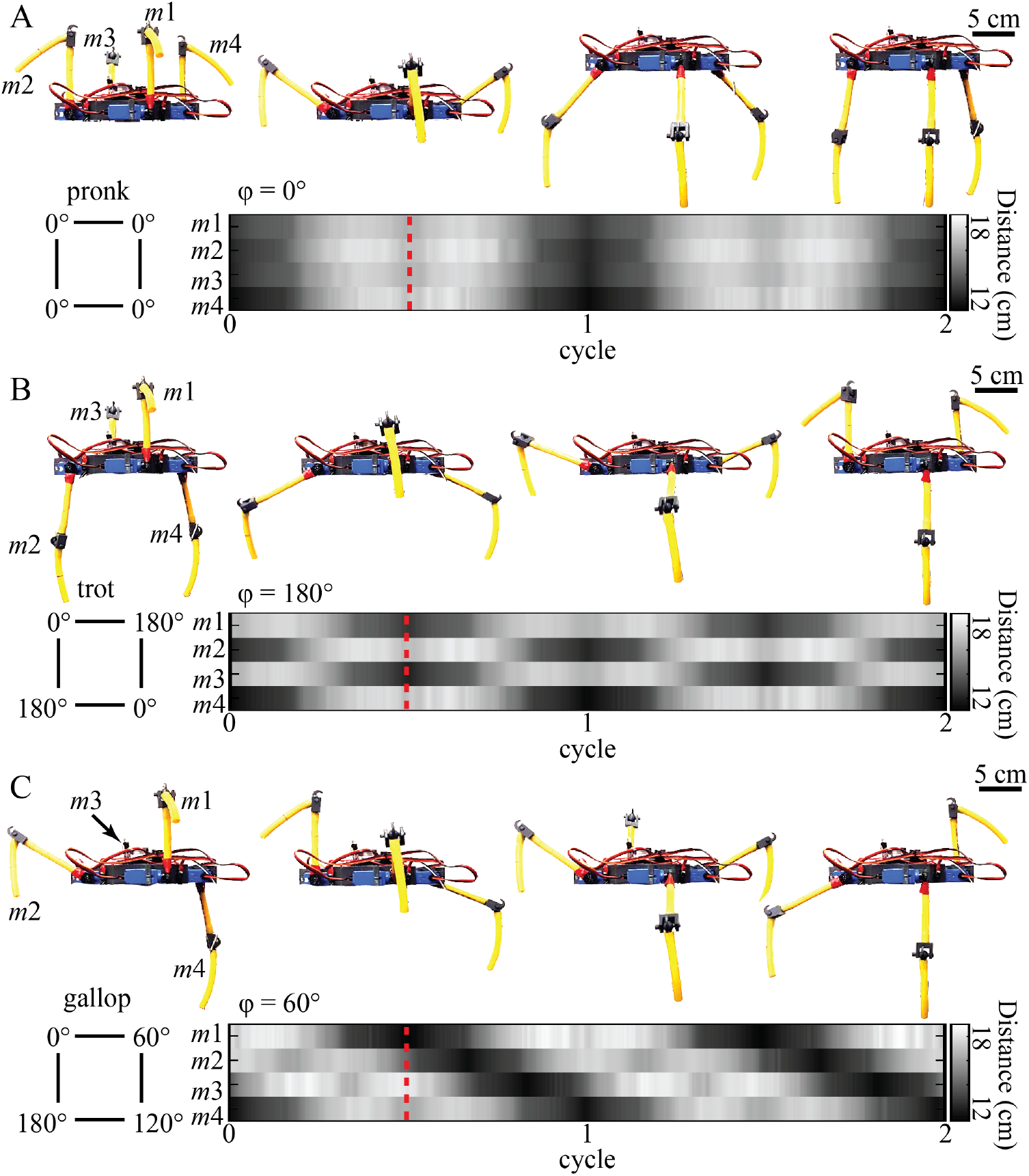
Quadriflagellate gaits prescribed to the robot. Distance from the center of geometry of the robot to the tip of each flagella was used as a proxy for the phase between adjacent flagella, labelled *m*1-4. (A) The pronk gait: zero phase difference (*φ*=0°) between adjacent flagella. (B) The trot gait: alternating pairs of flagella with a phase difference of half a gait cycle (*φ*=180°). (C) The gallop gait: adjacent flagella with a phase difference of a quarter of a gait cycle (*φ*=60°). Snapshots of the robot showing the flagella configurations during each gait over half a gait cycle. The dashed red line delineates half a gait cycle from the start of the recording. [Note to visualise the gaits fully the robot was not placed in fluid.]

For the movement of the robot in mineral oil (kinematic viscosity *µ/ρ* = 10 cm^2^/s), the Reynolds number (*Re*) 0.14 (*L* = 3.8 cm, *U* = 0.38 cm/s), while the oscillatory Reynolds number *Re*^osc^ was 0.20 (*L* = 3.8 cm, *ω* = 0.14 Hz). For the experiments conducted in glycerin (kinematic viscosity *µ/ρ* = 11.83 cm^2^/s), *Re* = 0.23 (*L* = 6.89 cm, *U* = 0.40 cm/s), and *Re*^osc^ = 0.55 (*L* = 6.89 cm, *ω* = 0.14 Hz).

### 2.4. Prescribing the swimming gait in the roboflagellate

We imposed three distinct gaits observed in quadriflagellate algae – the pronk, the trot, and the gallop. The different coordination patterns were achieved by prescribing the phase differences between adjacent appendages. The resulting gait sequences can be confirmed for an immobilised robot body, where the distance from each flagellum tip to the cell body was used as proxy for phase. In the pronk gait, all four appendages move simultaneously, without any phase difference (*φ* = 0°) between adjacent flagella (Figure 3A). The trot gait is defined by alternating pairs of flagella each of which is generating a pattern analogous to a breaststroke, with a phase difference of half a gait cycle (*φ*=180°) (Figure 3B). In the gallop gait, each appendage moves with a phase difference of a quarter-gait cycle relative to its neighbour (Figure 3C). The directionality (clockwise or counter-clockwise) of the gallop gait is determined by the phase difference (*φ*) between the first appendage (*m*_1_) and an adjacent appendage (*m*_2_ or *m*_4_). We tested the gallop gait in both a clockwise (*φ*=60°between *m*_1_ and *m*_2_) and counter-clockwise (*φ*=180°between *m*_1_ and *m*_2_) direction.

### 2.5. Motion tracking

Due to the of opaqueness of the oil, we attached LEDs to the robot’s body to enable motion tracking. Further LEDs were attached to the flagella. All LEDs were digitized using custom MATLAB algorithms. We approximated the center of geometry of the robot by averaging the position of the LEDs over time. Then, we used the tracks to determine the distance traversed by the robot in units of body lengths per beat cycle. A total of 9 trials were taken per gait, for each robot configuration. A trial was terminated either when the robot touches a boundary, or if the LEDs were no longer visible as the robot sediments over time (this is due to the foam trapping fluid and increasing in mass). Thus, each trial comprised 6 −10 cycles per gait. To visualize and confirm movement of the flagella during active swimming, we used glycerin as an alternative high viscosity fluid. However, because glycerin is not a dielectric fluid, wi-fi connectivity was interrupted and the circuits were negatively affected. To resolve this, we substituted our micro controller (Pro Trinket, Adafruit, product ID: 2000) and sealed the circuits with a gasket and a 3D printed cap.

## 3. Results and Discussion

### 3.1. The trot is the fastest gait in the algae

We identified the quadriflagellates as an ideal study group owing to their morphological diversity (in size, shape, aspect-ratio), and abundance in marine, terrestrial as well as freshwater habitats. A key trait distinguishing quadriflagellate genera is the arrangement or insertion of flagella around the anterior of the cell [21, 37]. Here we take advantage of this diversity to compare the swimming behaviour of three species (*Pyramimonas tetrarhynchus, Pyramimonas parkeae*, and *Carteria crucifera*) that employ three distinct gaits - respectively the pronk, the trot, and the gallop (Supplementary Video 1). We conjecture that inter-species differences in quadriflagellate swimming performance can be attributed to differences in gait alone - where the same basic stroke is applied to ensembles of appendages but according to distinct phase relationships.

Two of these algae belong to the genus *Pyramimonas*, a Prasinophyte algae belonging to an early diverging class which is thought to have given rise to the core Chlorophyte algae, comprising species with two, four, eight, or up to sixteen flagella [38, 5]. Four flagella of identical length and beat pattern emerge from an deep anterior groove or pit in the cell body. The third species, *C. crucifera*, is a Volvocalean flagellate that is closely related to the model biflagellate *Chlamydomonas*. Despite this phylogenetic divergence, all three species are similar in body size and flagellar morphology (approx ∼ 20 *µ*m in length and ∼15 *µ*m in width) and appear obovoid to cordate in side profile [39, 40].

In all three cases, cells swim smoothly flagella-first (puller-type) at speeds of 𝒪 (100) *µ*m*/s*. The translational motion is coupled to an axial rotation to produce swimming along helical trajectories [41]. Abrupt gait transitions can occur either spontaneously or when triggered by mechanical contact, during which the flagella are directed to the front of the cell in a so-called shock-response [42]. Cells can also reversibly stop and start swimming, when all or some of the flagella transiently cease to beat [18].

In all cases, free-swimming trajectories are superhelical, where small-scale swirls at the scale of single-cells are produced by the periodic flagellar oscillations. Three representative tracks, projected onto the focal plane, are shown in Figure 4 (A,C,E). Using the large-scale tracks, we estimated for each of the three gaits the displacement per cycle, including the cumulative displacement as a function of phase during the beat cycle (Figure 4G) as well as the mean forward progress per complete cycle (Figure 4H). Measured swimming speeds were 126 ± 24 *µm/s* for the pronk, 408 ± 46 *µm/s* for the trot, and 127 ± 25 *µm/s* for the gallop. Our results show that the trot gait is the fastest gait in the microalgae. Meanwhile the pronk and gallop gaits lead to comparable propulsion speeds.

**Figure 4.**
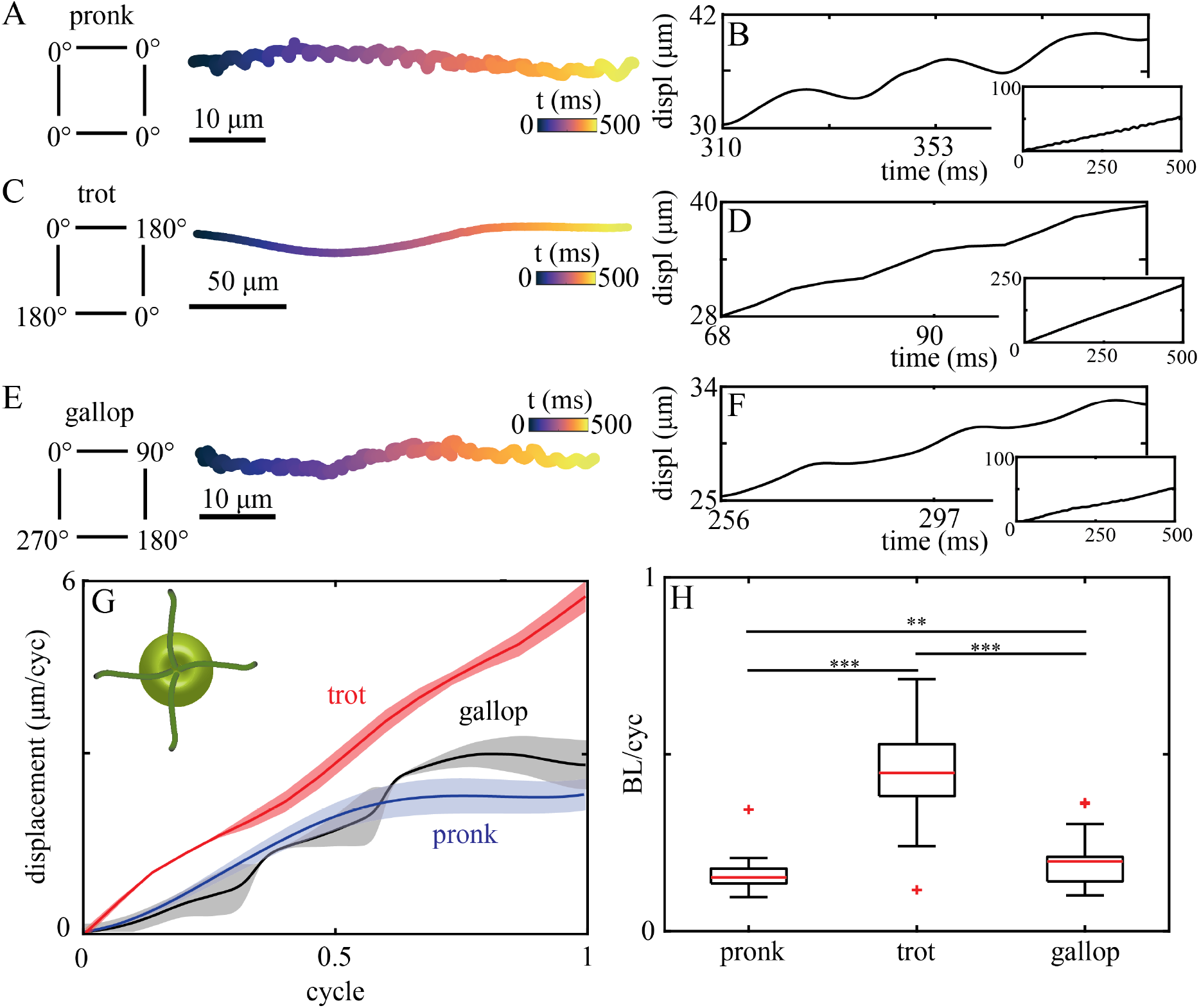
Gaits, kinematics, and hydrodynamic performance of quadriflagellate algae. All experiments were conducted in culture media - which had the same viscosity as water. For the pronking gait of *Pyramimonas tetrarhynchus*: (A) a sample (cell-centroid) trajectory colored by time, and (B) forward displacement over time for three cycles. Inset shows forward displacement over time of trajectory. For the trotting gait of *Pyramimonas parkeae*: (C) a sample (cell-centroid) trajectory colored by time, and (D) forward displacement over time for three cycles. Inset shows forward displacement over time of trajectory. For the galloping gait of *Carteria crucifera*: (E) a sample (cell-centroid) trajectory colored by time, and (F) forward displacement over time for three cycles. Inset shows forward displacement over time of trajectory. (G) Mean displacement within a gait cycle for all gaits - the pronk (blue line), trot (red line), and gallop (black line). Shaded areas correspond to the standard deviation. (H) Mean displacement computed in terms of body lengths per cycle, for each gait.

### 3.2. A hinged flagellum breaks time-reversal symmetry

We first confirmed that our system resides in a low-Reynolds number regime by attaching 3D-printed rigid (unhinged) flagella to the body (Supplementary Video 3). As expected, reciprocal strokes produced neglegible net swimming. The net displacement in the direction of movement after one complete cycle was 0.07 ± 0.05 *cm* (0.02 ± 0.01 BL) using the trot gait (Figure 5B).

**Figure 5.**
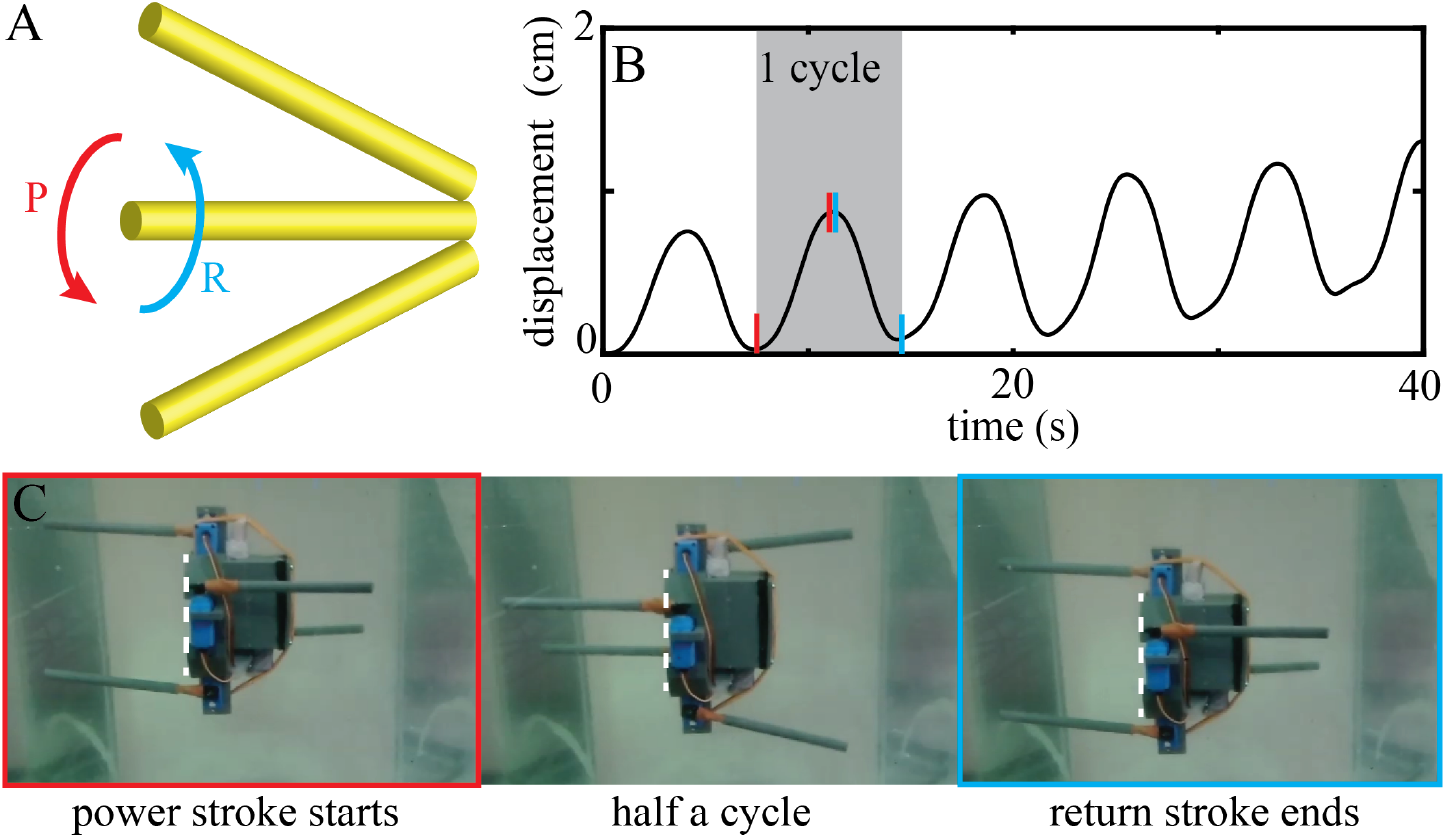
Kinematic reversibility confirms low-Reynolds regime. (A) One beat cycle of a single rigid flagellum moving back and forth. P: power stroke, R: recovery stroke. (B) Forward displacement traveled over time in mineral oil. Negligible net displacement per cycle with symmetric stroke pattern. (C) Snapshots of the robot during one gait cycle using rigid flagella. Left panel (outlined in red) shows the robot initiating a power stroke. Middle panel shows the robot during half a cycle. Right panel (outlined in blue) shows the end of the recovery stroke. [Note snapshots shown correspond to our alternative robot swimming in glycerin.]

With hinged flagella (Figure 2B), the robot became capable of net forward propulsion. Each gait cycle can be characterized by a power stroke during which the robot gains distance, and a recovery stroke during which it loses distance. We first set out to test the effect of flagella ‘waveform’ on swimming performance, this is expected to scale approximately with stroke amplitude [43, 44].

### 3.3. Flagellar undulation pattern affects swimming performance

We implemented two distinct flagellar undulation patterns - as defined by the maximal sweep range of the segments. For simplicity and to prevent axial rotation, we then reduced our quadriflagellate robot to a biflagellate robot, by removing one pair of flagella (Supplementary Video 4). The remaining pair of flagella was programmed to follow a breaststroke pattern (Figure 6A). We prescribed and compared the swimming performance for two different sets of motor angles for the proximal segment: i) [0°, 180°], and ii) [45°, 135°] (Figure 6A inset). The motion of the distal segment always follows passively, with the hinge breaking time-reversal symmetry. We tracked the flagella ‘waveform’ in the two cases and calculated the angles generated by each flagellum segment over time (from motor to joint and from joint to tip, Figure 6A). The two sweep amplitudes produced two distinct gaits *θ*_1_-*θ*_2_ shape space Figure 6B. A reduced sweep range results in a higher beat frequency (*ω* = 0.14 Hz for motor angles of [0°, 180°], and *ω* = 0.41 Hz for motor angles of [45°, 135°]). The rescaled displacement shows swimming performance increases with amplitude (Figure 6C). As predicted, the larger-amplitude breaststroke achieves a greater displacement after each gait cycle, consistent with the notion that non-inertial locomotion is dictated by simple geometric mechanics. Here, movement is kinematic, and net displacement is determined largely by the gait and its associated low-dimensional properties [45].

**Figure 6.**
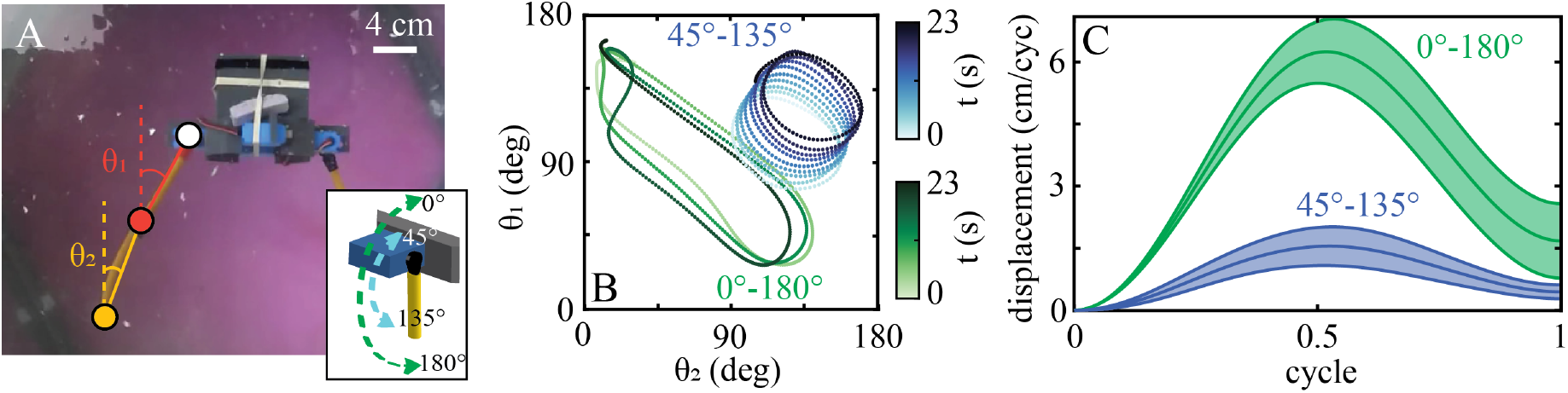
Swimming performance increases with stroke amplitude. (A) Quadriflagellate robot modified as a biflagellate robot, performing a breaststroke pattern with one pair of flagella. Angles *θ*_1_ and *θ*_2_ correspond to the angles generated by the flagella segment from the motor (white circle) to the joint (dark orange circle) and the segment from the joint (dark orange circle) to the tip (light orange circle). Inset shows variation of prescribed angles from 0°to 180°(green) and from 45°to 135°(blue). (B) *θ*_1_ as a function of *θ*_2_, colored by time. Green dots corresponds to angles from 0°to 180°. Blue dots corresponds to 45°to 135°. (C) Mean displacement as a function of a gait cycle. Green line corresponds to angles from 0°to 180°. Blue line corresponds to 45°to 135°. Shaded areas correspond to the standard deviation. These experiments were conducted in glycerin with the alternative robot, to ensure the flagella beat pattern can be tracked.

### 3.4. Roboflagellate swimming performance depends on gait and appendage placement

To test if swimming performance is dominated by gait or by other factors such as flagellar stiffness or shape dynamics, we prescribed the gaits exhibited by each algae species to our roboflagellates. We explored the effect of varying appendage phase coordination (gait) for two different configurations of four flagella, in which motors are positioned either in a parallel or a perpendicular orientation with respect to an identical body.

These configurations are inspired by naturally-occurring arrangements of basal bodies and flagella found in extant algal flagellates (Figure 7). All three species of algae studied here correspond to configuration A, in which the approximate plane of flagellar beating is perpendicular to the surface of the robot body. The main difference is that when viewed from the anterior of the cell, the four flagella are inserted with a clockwise twist or offset for *Carteria*, but an anticlockwise offset for *Pyramimonas* [37]. Algal species reported to exhibit configuration B [37] were not available in culture and were not represented in the present study. Appendage coordination was prescribed in the robot by specifying the phase differences between flagella, to produce each of the three gaits: pronk, trot, or gallop, as previously described Figure 3.

**Figure 7.**
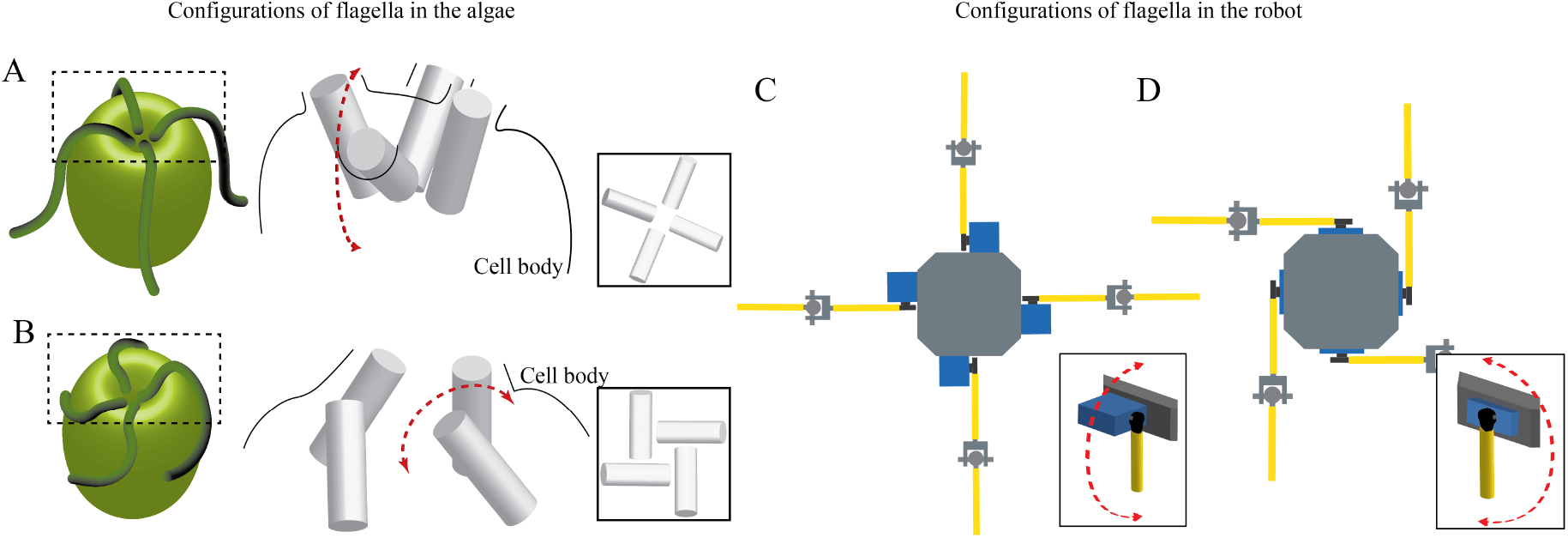
Modelling appendage placement. (A) Illustration of two configurations of flagella and basal bodies that are found in quadriflagellates [37]. The flagella emerge from basal bodies (cylinders) that are oriented largely perpendicular (A), or parallel (B) to the cell body. Insets show anterior views (A: cruciate arrangement, B: turbine or windmill-like). Double-arrow indicates the approximate beat plane of the individual flagella. Similarly, two roboflagellate designs are presented. Motors and attached ‘flagella’ are oriented perpendicular (C) or parallel (D) to the central body. Again, double-arrow indicates oscillation plane.

For the perpendicular configuration, example trajectories as well as the cumulative forward displacement over time for each gait as shown in Figures 8(B)-(C), (E)-(J). We also analyzed the detailed within-cycle dynamics for each gait (Supplementary Video 5). The pronk and both the clockwise (CW) gallop and counterclockwise (CCW) gaits produce significant forward displacement during the power stroke (up to 5.7 cm for the pronk, 4 cm and 2 cm for the CW and CCW gallops respectively after 1/2 a gait cycle), but also produce a significant backward displacement during the recovery stroke, generating overall small displacement from cycle to cycle (0.33±0.04 BL/cyc, 0.16±0.05 BL/cyc, and 0.15±0.08 BL/cyc for the pronk, the CW gallop, and CCW gallop respectively). On the other hand, while the trot does not achieve a greater displacement (only 2.3 cm after 1/2 a gait cycle) than the pronk or gallop during the power stroke, it loses a much smaller distance during the recovery stroke. This is because while one pair flagella is moving towards the body and consequently producing backward motion, the other pair of flagella moves away from the body so as to resist this motion. This can also be observed in the trajectories, where the pronk and gallop gait shows backward motion, unlike the trot gait. Due to this, of the three gaits investigated the robot achieves the greatest hydrodynamic performance (0.6±0.08 BL/cyc) using the trot gait (Figure 8K), just as in the algae.

**Figure 8.**
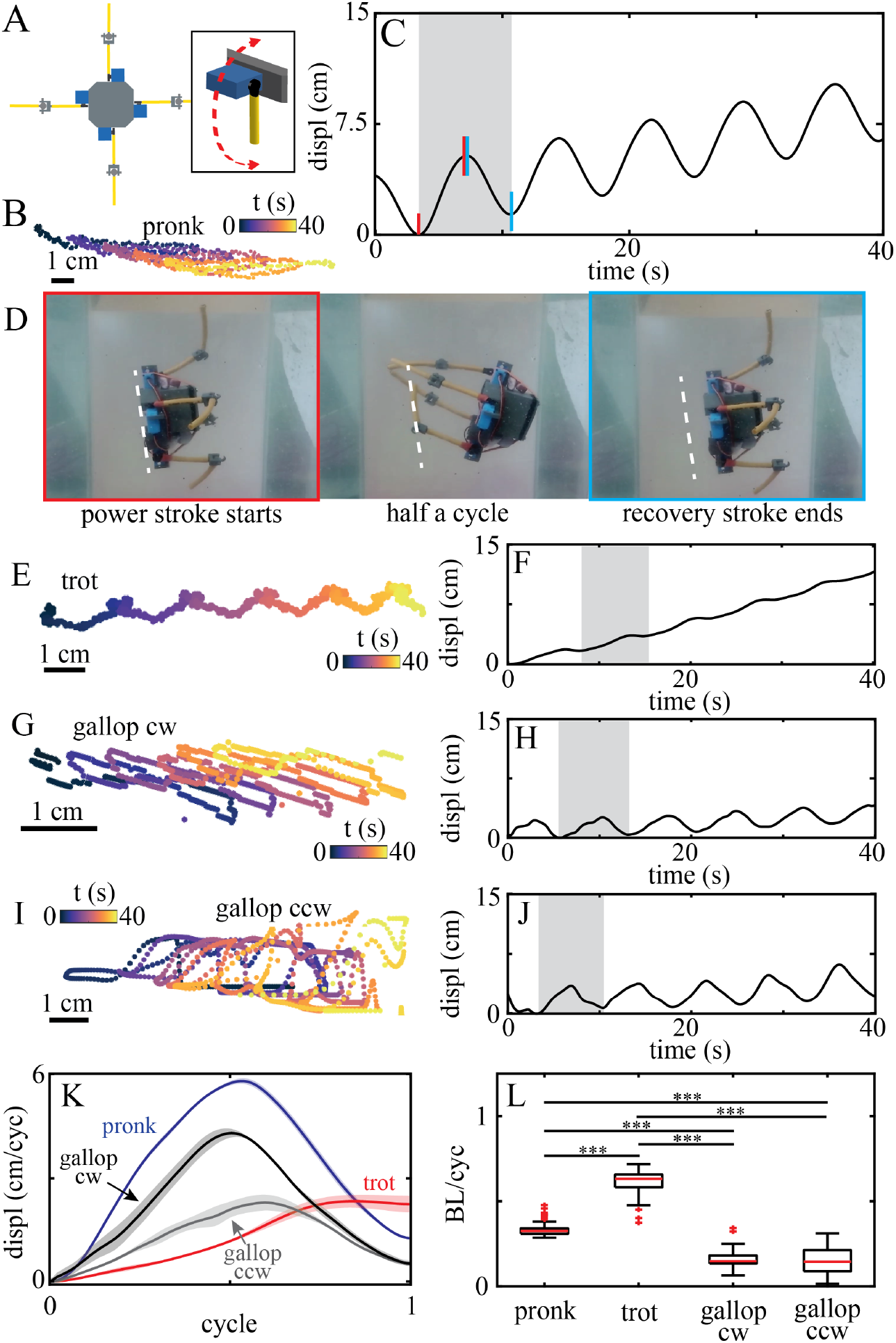
Swimming gait kinematics and performance for robot with flagella in perpendicular orientation. (A) Diagram of robot with motors oriented perpendicular to the body. Inset illustrates beating plane. For the pronk gait, (B) shows a sample trajectory of the robot, colored by time (5 cycles), and (C) the forward displacement traveled over time. For one gait cycle, red vertical lines highlight power stroke, and blue vertical lines highlight return stroke. (D) Snapshots of the robot during one cycle of the pronk gait. Left panel (outlined in red) shows the robot initiating a power stroke. Middle panel shows the robot during half a cycle. Right panel (outlined in blue) shows the robot completing the recovery stroke. (Arrow: swimming direction.) Trajectory of the robot during the trot gait, colored by time (5 cycles) (E), and forward displacement traveled over time of the robot during the trot gait (F). Trajectory of the robot during the clockwise gallop gait, colored by time (5 cycles) (G), and forward displacement traveled over time of the robot during the trot gait (H). Trajectory of the robot during the counter-clockwise gallop gait, colored by time (5 cycles) (I), and forward displacement traveled over time of the robot during the trot gait (J). (K) Mean displacement over a gait cycle for all gaits - the pronk (blue line), trot (red line), clockwise gallop (black line), and counter-clockwise gallop (grey line). Shaded areas correspond to the standard deviation. (L) Body length per cycle as a function of swimming gait.

For the parallel configuration (Supplementary Video 6), example trajectories as well as the forward displacement over time for each gait can be seen in Figure 9(B)-(C), (E)-(J). Similar to the perpendicular configuration, the pronk gait allows the robot to gain a significant amount of distance during the power stroke (up to 5 cm after 1/2 a gait cycle) but also lose a significant amount of distance during the recovery stroke, generating little net displacement from cycle to cycle. The gallop gait in the counterclockwise displays a similar oscillatory pattern, however there is a discrepancy between the counterclockwise and clockwise gallops (5.3 cm after 1/2 a gait cycle for the CW gallop, but only 1 cm for the CCW gallop). This is likely due to rotation-translation coupling in the second configuration (in which the flagella are inserted in the CCW sense), generating significant motion laterally and causing axial rotation of the robot. Similar to the perpendicular robot, the trot gait gains less distance during the power stroke (only 1.5 cm after 1/2 a gait cycle) and loses more distance during the recovery stroke, relative to the perpendicular configuration. The phasing between appendages in the trot gait again aids the robot in traversing a greater distance from cycle to cycle than the pronk (0.15±0.4 BL/cyc), and also greater than the average of the CW and CCW gallop gaits (0.15±0.9 BL/cyc). (We assume that by symmetry, this average between the two chiralities should cancel any rotational effects.) Thus, the trot remains a hydrodynamically effective gait for the parallel robot (0.26 ±0.08 BL/cyc) (Figure 9K).

**Figure 9.**
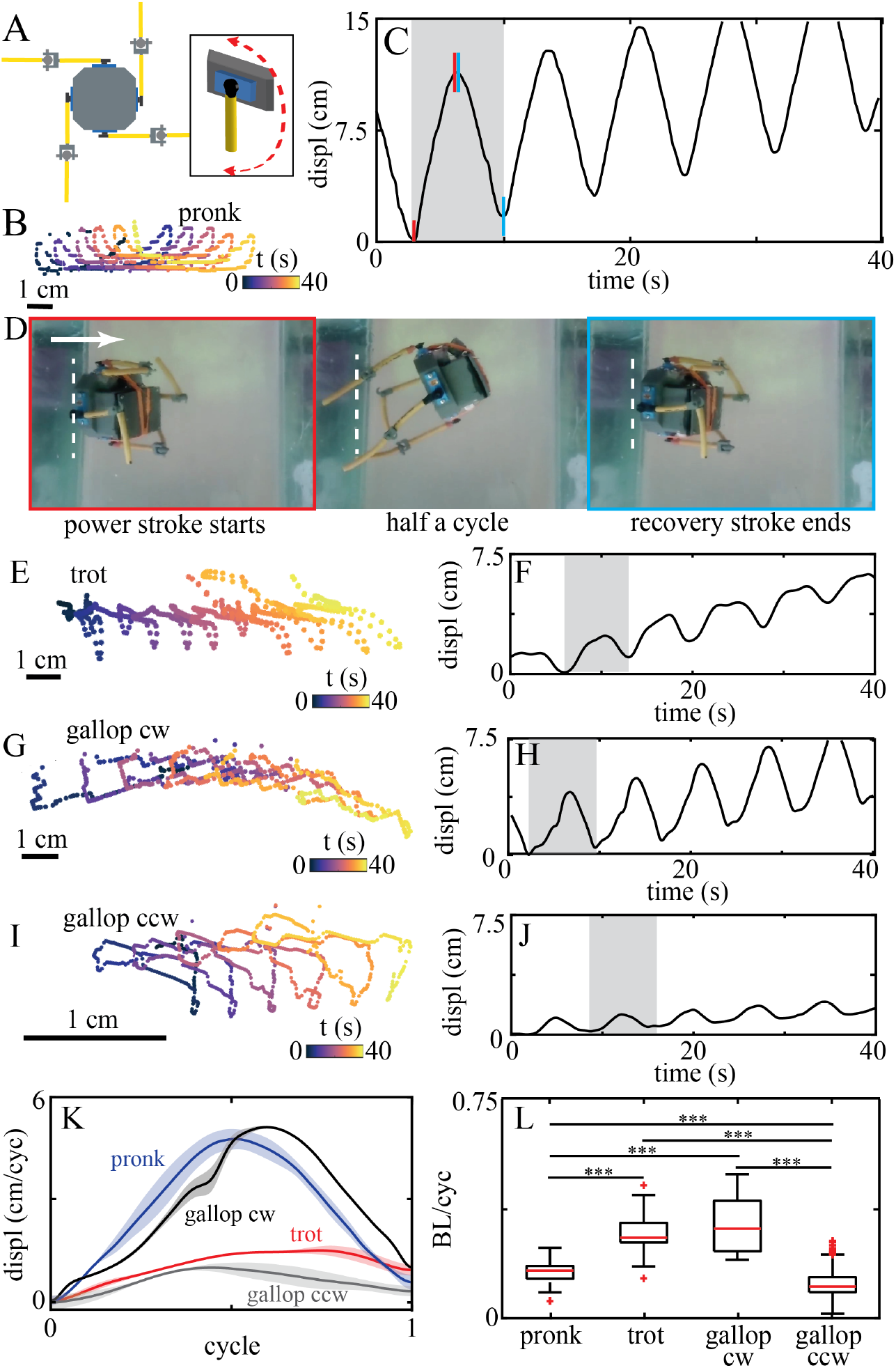
Swimming gait kinematics and performance for robot with flagella in parallel orientation. (A) Diagram of robot with motors oriented perpendicular to the body. Inset illustrates beating plane. For the pronk gait, (B) shows a sample trajectory of the robot, colored by time (5 cycles), and (C) the forward displacement traveled over time. For one gait cycle, red vertical lines highlight power stroke, and blue vertical lines highlight return stroke. (D) Snapshots of the robot during one cycle of the pronk gait. Left panel (outlined in red) shows the robot initiating a power stroke. Middle panel shows the robot during half a cycle. Right panel (outlined in blue) shows the robot completing the recovery stroke. (Arrow: swimming direction.) Trajectory of the robot during the trot gait, colored by time (5 cycles) (E), and forward displacement traveled over time of the robot during the trot gait (F). Trajectory of the robot during the clockwise gallop gait, colored by time (5 cycles) (G), and forward displacement traveled over time of the robot during the trot gait (H). Trajectory of the robot during the counter-clockwise gallop gait, colored by time (5 cycles) (I), and forward displacement traveled over time of the robot during the trot gait (J). (K) Mean displacement over a gait cycle for all gaits - the pronk (blue line), trot (red line), clockwise gallop (black line), and counter-clockwise gallop (grey line). Shaded areas correspond to the standard deviation. (L) Body length per cycle as a function of swimming gait.

We conclude that the swimming performance of the roboflagellate is highly sensitive to both gait and flagellar orientation (which defines the principal beat plane) of the flagella. It is possible that the organisms can access different regimes by controlling the 3D beat plane of their flagella, and that divergent flagellar placement evolved in different species as a result of different environmental selection pressures. In flagellates such as *Volvox*, nearby basal bodies (from which the flagella emerge) have rotated 90 degrees compared to the ancestral configuration found in the unicellular *Chlamydomonas*, likely to facilitate coordinated flagellar beating as an intact colony [5, 46].

### 3.5. Speed of roboflagellate is comparable to that of real algae

The above results show that a change in flagellar configuration can significantly change the performance of a given swimming gait. Focusing only on the trot, we note that the trot gait yielded the highest hydrodynamic performance for the algae and for the perpendicular robot, (Figure 10).

**Figure 10.**
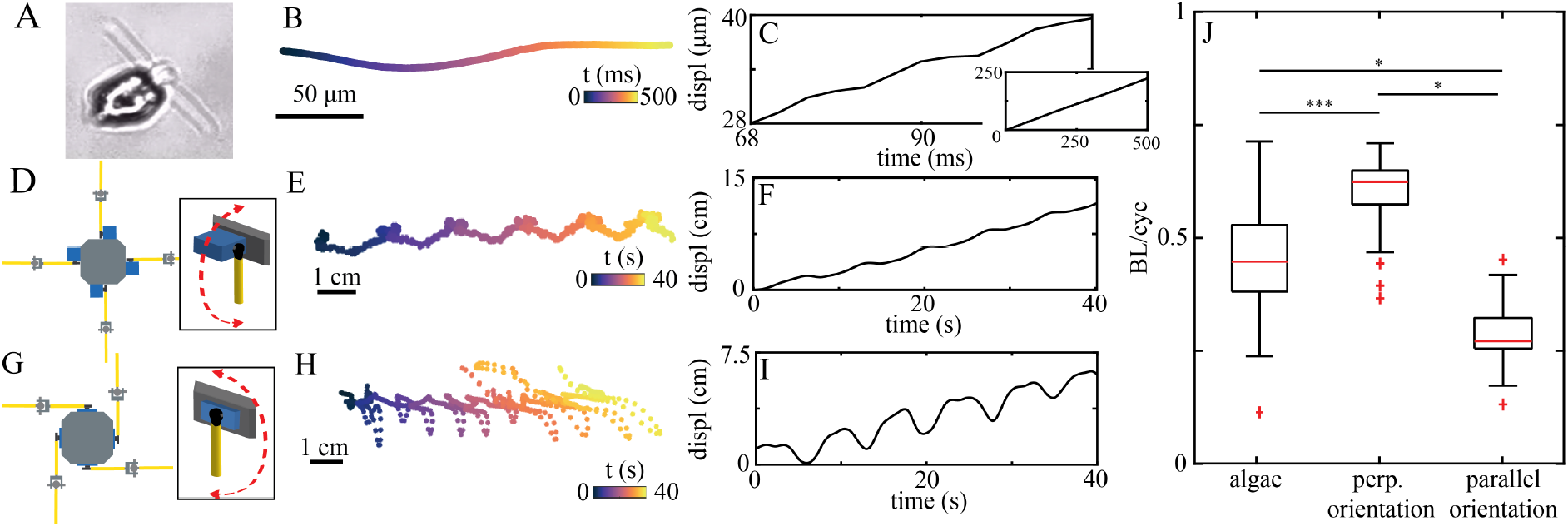
Comparing the trot gait in the algae and robot. (A) The alga *Pyramimonas parkeae* swimming using the trot gait. (B) Trajectory of *P. parkeae*, colored by time. (C) Forward displacement traveled over time by *P. parkeae*. (D) Diagram of robot with motors oriented perpendicular to the body. Inset illustrates beating plane. (E) Trajectory of the robot with perpendicular configuration using the trot gait, colored by time. (F) Forward displacement traveled over time of the robot with perpendicular configuration using the trot gait. (G) Diagram of robot with motors oriented parallel to the body. Inset illustrates beating plane. (H) Trajectory of the robot with parallel configuration using the trot gait, colored by time. (I) Forward displacement traveled over time of the robot with parallel configuration using the trot gait. (J) Body length per cycle for the trot gait for the algae, the perpendicular configuration, and the parallel configuration.

In both robot configurations, significant axial rotation and lateral movement was observed in the free-swimming trajectories (Figure 10E,H) showing that our robotic models do not swim as smoothly as their algal counterparts (Figure 10B). This suggests that the algal cytoskeleton could play a role in gait stabilization. The cumulative displacement over time for a trotting cell and our perpendicular robot are comparable (Figure 10C,F). Meanwhile the parallel configuration displays larger amplitude oscillations in which a greater distance gained during each the power stroke is negated during the subsequent recovery stroke (Figure 10I). This is likely due to three-dimensional effects as mentioned above. In all, we find that the performance of the algae and both roboflagellate configurations are comparable in absolute terms, as measured in terms of body lengths per stroke cycle. This agreement is surprising as we did not precisely match the dimensions of our robots to that of the algal cell, and unlike the algal flagella the robot ‘flagella’ were not capable of active bending - being comprised only of rigid tubing and a 3D-printed hinge.

## 4. Conclusion and future work

Microscopic organisms have evolved to harness many different ways of swimming at low-Reynolds number. Despite their apparent simplicity and lack of centralized control, many species of single-celled algal multiflagellates can perform robust free-swimming gaits analogous to animal gaits, which require specific temporal ordering of a small network of flagella [18]. Since structural and genetic information about many of the species of interest is lacking, we turned to robophysical modelling to understand how motility and gait are controlled in these unicellular microswimmers.

Robotics approaches at the macroscale have long been used successfully to explore and validate fluid dynamical theories of self-propulsion [47, 48]. Scenarios can be tested in robots that may not be possible in the live organism [49, 50]. Dynamically-scaled robotic models of microswimmers operating in viscous media have been used to mimic bacterial swimming [51], to examine flows induced by bundles of rotating flagella [52], and to reveal the role of elasticity [53], or to investigate metachronal actuation of rigid appendages [54].

Here, we created for the first time a novel self-powered, untethered robot (no external forces or torques) modelled upon quadriflagellate algae, which could not only self-propel at low-Reynolds number but also recapitulate gait-dependent differences in swimming performance that were observed in different species of microalgae. These results reveal that differences in phase coordination of propulsive appendages alone has a significant impact on hydrodynamic performance. The orientation of moving appendages on the propelling body also influences net propulsive speed. In the perpendicular configuration that best matches the algae, the trot gait is consistently faster than either the pronk or gallop gait, and that the net displacement achieved by the robot in terms of body lengths per cycle is similar in absolute terms to the algae. Thus our dynamically-scaled robophysical model is a good model of the biological microswimmer.

Our work raises open questions about why the different quadriflagellate species have distinct motility repertoires in the first place. Freely-locomoting organisms at all scales, switch dynamically between multiple gaits, [55, 56], e.g. to escape predation. Even bacteria motility exhibits strong heterogeneity across species [3]. While marine *Pyramimonas* species exhibit sporadic bursts of fast activity with extended quiescent phases [42], freshwater alga *Carteria* (closely related to *Chlamydomonas*) and other *Volvocalean* algae do not show such rest periods [18]. More generally, ciliary strokes that are optimised for swimming may not be optimised for other tasks such as feeding or fluid pumping, and vice versa [57, 58]. Distinct gaits are unlikely to have evolved to achieve ever-faster swimming, but rather reflects a more nuanced relationship between the organism’s metabolic requirements and its habitat.

We conjecture that differences in gait confers an evolutionary advantage even at the microscale. Of the three algae studied here, two (*P. tetrarhynchus, C. carteria*) are freshwater species and one is a marine species (*P. parkeae*). *P. tetrarhynchus* (type species) was originally isolated from a peaty pool and cultured in a biphasic soil medium [39]. *C. crucifera* is also a freshwater species that forms surface associations with leaves and other decaying material. In contrast *P. parkeae* is most abundant in Acrtic surface water and in tidal rock pools, where it can access sufficient sunlight for photosynthesis. *P. parkeae* also exhibits a unique diurnal vertical settling behaviour [59]. The latter behaviour, along with phototaxis, accentuates the requirement for vigorous swimming and hence the fast trot gait. Field data has shown that marine *Pyramimonas* routinely blooms in and around sea ice, where the unique polar environment (extreme fluctuations in temperature, light, salinity etc) is associated with a highly heterogeneous distribution of different *Pyramimonas* species even within the same water column [60]. The habitats of these algae may therefore be a key evolutionary driver leading to significant diversification of gait, even across species with apparently convergent morphology and size [61, 62]. Further experiments using both lab strains and wild isolates, controlling more precisely for culturing medium, are need to test this hypothesis. In parallel, we will use roboflagellates to explore mix-mode propulsion strategies and unsteady effects, such as nutrient dispersal.

We highlight two limitations of the current design. The first concerns boundary and finite-size effects, particularly due to fluid-structure interactions between moving appendages and the bounding tank, and between different parts of the robot. The presence of no-slip boundaries will alter the flow fields around a beating appendage, and change propulsion efficiency [63]. The rigid insertion of the robot flagella around the central body likely introduced an additional (unwanted) rotational movement. Second, the current robot relies on a simple 2-link flagellum facilitated by a rigid 3D printed joint which has a very limited number of degrees of freedom. The rigid joints have limited ability to resist torsion - which may be gait-dependent. The compliance of real (eukaryotic) flagella and cilia, which deform actively by distributed motor elements, also improves propulsive force generation and efficacy. These organelles can actively maintain their shape even when subject to significant hydrodynamic forces. In future work we will resolve these limitations with improved roboflagellate designs, in parallel with hydrodynamic simulations and modelling to understand gait optimisation.

In conclusion, we have presented a macroscopic robot capable of self-propulsion at low-Reynolds number, and used this to model aspects of microorganism swimming behaviour. We have applied this physical model to different permutations of gait coordination patterns and flagellar placement and their influence on hydrodynamic swimming performance. This approach has transformative potential for testing new hypotheses relating to low Reynolds number self-propulsion in other small-scale biological systems, such as mechanisms of gait selection and stimulus-dependent steering [16]. These insights could have profound implications for how morphological computation may be achieved in aneural or early nervous systems. From a technological perspective, these diverse propulsion strategies can provide unique, innovative solutions to the formidable challenge of navigating viscous fluids.

## Acknowledgements

The authors are grateful for funding from Physics of Living Systems Student Research Network to KD and DIG, NSF Simons Southeast Center for Mathematics and Biology (SCMB) to KD, the Dunn Family Professorship (DIG), and from the European Research Council (ERC) under the European Union’s Horizon 2020 Research and Innovation Programme (grant agreement No 853560 to KYW). We also acknowledge the 2018 MBL Physiology Course (Woods Hole, MA) during which this project was conceived.

## Notes

### Competing Interest Statement

The authors have declared no competing interest.

